# Somatic Mutations Predict Acute Myeloid Leukemia Years Before Diagnosis

**DOI:** 10.1101/237941

**Authors:** Pinkal Desai, Nuria Mencia-Trinchant, Oleksandr Savenkov, Michael S. Simon, Gloria Cheang, Sangmin Lee, Michael Samuels, Ellen K Ritchie, Monica L. Guzman, Karla V Ballman, Gail J. Roboz, Duane C. Hassane

## Abstract

**Background:** Somatic mutations observed in clonal hematopoiesis are associated with increased age and risk of hematological malignancies. However, the limited number of acute myeloid leukemia (AML) patients in studies of clonal hematopoiesis thus far has precluded determination of the spectrum of mutations leading to AML and their impact on risk and time to diagnosis.

**Methods:** The relationship between baseline mutations and subsequent AML was determined in a case-control study design. 212 women eventually diagnosed with AML (median time: 9.8 years) were identified from the Women’s Health Initiative cohort alongside 212 matched AML-free controls. Deep sequencing of 67 genes was performed on DNA isolated from peripheral blood to detect subclonal mutations.

**Results:** The presence of any mutation was associated with increased odds of eventual AML (odds ratio [OR] 4.0; 95% confidence interval [CI], 2.5-6.3). These odds were further elevated with mutations in IDH1/2 (OR 8.4; 95% CI, 1.4-51.9), TP53 (OR 54.2; 95% CI, 2.9-1017.7), or spliceosome genes (OR 5.6; 95% CI, 1.5-20.6). Eventual AML diagnosis occurred in all or most participants with mutations in TP53 (N=23/23) or IDH1/2 (N=15/16). Mutations indicated sooner AML diagnosis (median 8.0 vs. 11.9 years; P < 0.001) with TP53 mutations demonstrating increased odds of AML within 5 years (OR 3.6; 95% CI, 1.4-9.1).

**Conclusions:** Mutations are present in peripheral blood of AML patients a decade prior to AML diagnosis with mutations in TP53 producing especially rapid presentation. Strategies for monitoring of high-risk populations are needed and clinical trials of potential early interventions can be considered.

## INTRODUCTION

The pathogenesis of acute myeloid leukemia (AML) is characterized by serial acquisition of somatic mutations and several genes are recurrently mutated in AML (1–3). However, it is not known when such mutations appear prior to the development of overt disease, how they evolve, and the specific risk associated with each one. Furthermore, the acquisition of AML-associated mutations has also been found in normal aging, with approximately 10% of persons greater than 65 years of age (4–7) having so-called “clonal hematopoiesis of indeterminate potential” (CHIP). The presence of CHIP is associated with an elevated risk of hematologic malignancies and cardiovascular disease. However, studies of CHIP to date have included very few subjects who subsequently developed AML. Jaiswal et al. performed whole exome sequencing on peripheral blood samples from 17,182 individuals (5). Mutations in genes associated with hematologic cancers were found in 5.6% of subjects aged 60-69 years and in up to 18.4% of those aged >90 years. Only 16 hematologic cancers were reported in this group, of which 6 were AML cases. In a study by Genovese et al, DNA sequencing was performed on peripheral blood from 12,380 Swedish individuals (mean age 55 years), with clonal mutations detectable most commonly in DNMT3A, ASXL1 and TET2. Clonal hematopoiesis was strongly associated with increased risk of hematologic cancer (hazard ratio, 12.9; 95% C.I., 5.8 < 28.7), but there were only 12 AML cases reported in the study (4). The objective of the present work was to investigate whether AML-associated mutations could be detected years prior to the onset of AML and to determine whether specific mutations, allele burdens, or patterns of coexisting mutations would affect the risk and time-to-diagnosis of AML. To this end, we performed deep sequencing of serially collected peripheral blood samples obtained from 212 women a median of 9.8 years prior to their diagnosis of AML, along with 212 matched controls.

## METHODOLOGY

### Study population and samples

The Women’s Health Initiative (WHI) was a large, prospective clinical trial and observational study initiated by the U.S. National Institutes of Health (NIH) in 1991 and consisted of three clinical trials designed to assess impact of hormonal therapy on postmenopausal health issues in women. More than 160,000 women were eventually enrolled in one or more of four clinical trials (CT group) or an observational study cohort (OS group) in 40 U.S. clinical centers from October 1, 1993 through December 31, 1998. Follow-up continued from study initiation until planned termination on March 2005. Thereafter, participants providing re-consent were followed for an average of 10.8 years (SD 3.3 years), with data collection updated through September 2012 (8). The participants in the CT group were followed at baseline and years 1, 3 and 6, with samples collected at WHI baseline, year 1 and 3 during these follow up visits. The participants in the OS group were followed at baseline and year 3,with samples collected similarly during these visits (8, 9). Detailed clinical data regarding medical history, medications, and complete blood counts were available at baseline assessment. New diagnoses were updated in follow up, with central confirmation of all new cancer diagnoses.

#### Identification of cases

In the WHI cohort, 212 study participants eventually developed pathologically confirmed AML. Of these, baseline peripheral blood (PB) DNA was available and passed quality control in 189 participants; these were identified as cases to be included in the final analyses. Additional follow up samples for 132 cases were available at 1 year and/or 3 years after baseline, all prior to the diagnosis of AML. Exclusion criteria for cases included known diagnosis of any myeloid disorder, including AML, prior to WHI baseline evaluation. Participants who had a diagnosis of AML within 6 months of WHI baseline assessment were also excluded.

#### Controls

Matched controls (N=212) were selected from participants who were confirmed not to have a diagnosis of AML while being followed on the WHI study. Exclusion criteria included concurrent or history of prior myeloid disorder, including AML, at WHI baseline. Controls were matched to cases by age at baseline, WHI component (clinical trial vs. observational study), and history of non-myeloid cancers at baseline. In addition to these criteria, controls were also matched by type and timing of any cancers that occurred in cases after WHI baseline, but before the diagnosis of AML. For example, if a clinical trial participant with AML had prior breast cancer and was age 60 at baseline, a corresponding 60-year-old control participant with prior breast cancer history was selected who never developed subsequent AML. Similarly, if a participant with AML (case) had breast cancer diagnosed after her year 1 PB draw, she was matched to a control who also had breast cancer diagnosed after her year 1 blood draw, but did not develop AML. Matching was done in a time forward manner to ensure that each control had as much control time as its matched case (10). For example, a participant who developed AML two years after randomization on the WHI protocol would be matched with a control with at least two years of follow-up. Of 212 controls, PB was available and passed quality standard in 183 controls at WHI baseline and these were included in the analyses. Additional follow up samples for 128 controls were available at 1 year and/or 3 years after baseline.

### Statistical Analysis

Baseline characteristics of AML cases and matched controls were compared with the use of the two-sample t-test for continuous variables and the Fisher’s exact test for categorical variables. Among the 189 cases, participants with baseline precursor mutations were compared to participants without precursor mutations with regard to demographic characteristics and baseline hematological characteristics (i.e. WBC count and differential counts, hemoglobin value and platelet count). The relationship between specific precursor mutations and AML development were estimated by exact odds ratios (OR) and adjusted ORs were obtained from penalized-likelihood logistic regression (11). OR and adjusted ORs are presented with their associated 95% confidence interval (95% CI). Multivariable penalized-likelihood logistic regression analysis was performed to assess the independent effect of demographic and prognostic factors of interest on precursor mutation status. Collinearity between predictors in the models was evaluated prior to the formulation of the final multivariable models. Time to development of AML was estimated with a Kaplan-Meier estimator. Differences between groups based on mutational status were evaluated with a log-rank test. Significant differences in variant allele fraction were determined in serial sampling using Fisher’s Exact Test based on the count of supporting alternate and reference reads for each sample at a mutated site. All p-values were two-sided with statistical significance set a priori at the 0.05 level. Ninety-five percent confidence intervals (95 % C.I.) were calculated to assess the precision of the obtained estimates. All statistical analyses were performed with the use of R software (version 3.4.0).

### Targeted Exome Sequencing

Genomic DNA was provided by WHI in a blinded manner, in which case-control status and clinical covariates were revealed only after variant calling was completed. Library generation and amplification were performed using a low error rate Hi-Fi DNA polymerase according to the Kapa HyperPrep protocol (Kapa Biosystems). Dual sample indexing, rather than single indexing, of libraries was performed to minimize signal spread errors arising from misidentification of multiplexed samples (12). Targeted sequencing using a panel of 67 recurrently mutated genes in hematological malignancies was performed using a custom capture probes (Nimblegen) to a median coverage of 2000x for both AML cases and controls (Figure S1). Variant analysis was performed following rigorous quality control and filtration of low quality sequence information. To identify somatic variants, filtration based on population allele frequency data was applied so as to enrich somatic variants that are not likely inherited. To this end, variants were classified as probable somatic if exhibiting a dbSNP v142 or ExAC adjusted population allele frequency <= 0.25% or a median VAF in the cohort < 40%. Only mutations present at >1% VAF were evaluated for association with AML development and time-to-AML. Further details are provided in the Supplemental Methods section.

## RESULTS

### Clonal mutations in peripheral blood are evident years prior to the diagnosis of AML

At a median of 9.8 years prior to the diagnosis of AML, cases were more likely to harbor mutations than controls (OR 4.0, 95% C.I. 2.5 – 6.3, P < 0.001). The most common mutations identified above 1% variant allele fraction (VAF) included DNMT3A (37.6% cases vs. 19.1% controls), TET2 (25.4% cases vs. 6.0% controls), TP53 (12.2% cases vs. 0% controls), SRSF2 (6.8% cases vs. 0.5% controls), IDH2 (6.3% cases vs. 0.5% controls), SF3B1 (5.8% cases vs. 0.5% controls), JAK2 (5.8% cases vs. 1.1% controls), and ASXL1 (3.2% cases vs. 4.4% controls; Fig. 1A and Figure S2). In aggregate, spliceosome mutations as a group (SF3B1, SRSF2, and U2AF1) were identified in 14.2% of cases (N=27/189) vs. 1. 6% of controls (N=3/183). Similarly, IDH mutations as a group (IDH1 and IDH2) were identified in 7.9% of cases (N=15/189) vs. 0.5% of controls (N=1/183). There was no association between the presence of any mutation and abnormal hemoglobin, white blood cell (WBC) count and/or platelet level (Supplemental Table 1).

**Figure 1.**
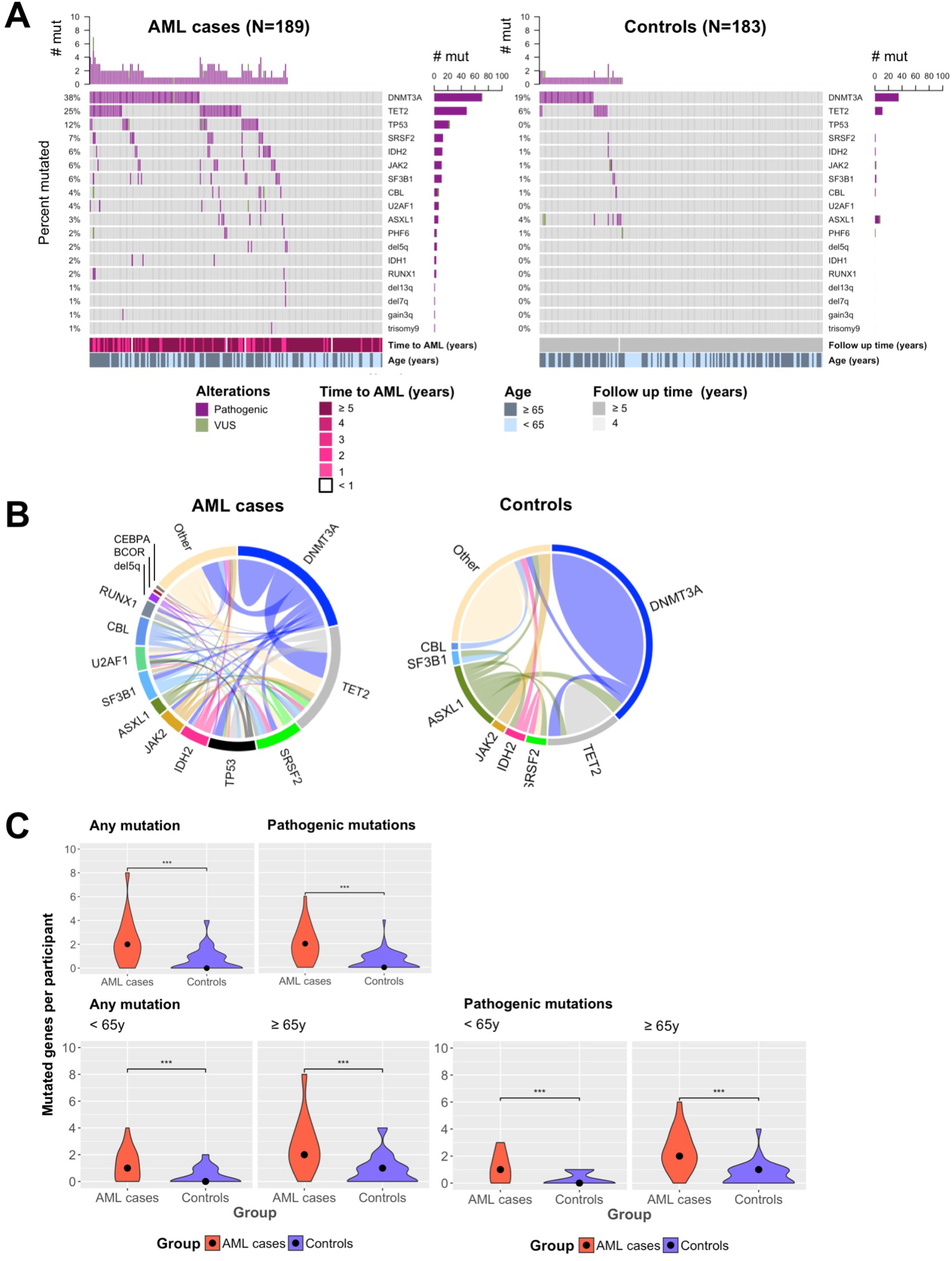
Spectrum of mutations seen years at baseline prior to the diagnosis of AML alongside matched controls. (A) OncoPrint summarizing mutated genes AML cases (N=189; left) vs. controls (N=183; right). For each gene (rows), pathogenic mutations (purple) are indicated vs. variants of unknown significance (VUS; green). Vertical barplots (right of each OncoPrint) indicate the number of participants in whom the gene is mutated (# mut) in each group. Horizontal barplots (top of each OncoPrint) indicate the number of mutated genes (# mut) in each participant. For the AML controls, the time to AML and age (≥ 65 years vs. ≤ 65 years) are shown. For the controls, the time of follow up and age (≥ 65 years vs. ≤ 65 years) are shown. (B) Chord plot indicates co-mutations between genes in AML cases alongside controls. The arc length is directly proportional to the percent of instances of co-mutation. Each color represents a different gene. “Other” includes genes not recurrently mutated in AML. (C) Violin plot indicating the distribution of the number of mutated genes per participant in AML cases (red) vs. controls (blue). Top row shows all mutations vs. pathogenic mutations. Bottom row further stratifies differences in mutation number into age groups (< 65 years vs. ≥ 65 years). *** P < 0.001, Wilcoxon test.

AML cases overall demonstrated greater clonal complexity than controls, with 55.6% of the AML cases harboring co-mutations, compared to 17.6% of controls (OR 5.2, 95% C.I. 2.5-11.7, P < 0.001). As shown in Fig. 1B, the most common co-mutations present in AML cases were DNMT3A with TET2, DNMT3A with TP53, DNMT3A with SRSF2, TET2 with SRSF2, and IDH2 with SRSF2. Among controls, mutations were generally present individually, and the only common co-mutation was DNMT3A with TET2. Mutations categorized as pathogenic (see Supplemental Methods) were also significantly increased in AML cases vs. controls (90% vs. 84%, respectively; P < 0.001). This was observed for both younger (< 65 yrs.) and older participants (≥ 65 yrs.) (Fig. 1C).

### The presence of somatic mutations at baseline assessment was associated with significantly increased odds of developing AML

Having a mutation at baseline assessment was associated with statistically increased odds of developing AML (OR 4.0, 95% C.I. 2.5-6.3, P < 0.001) (Table 1A). This finding was independent of age: OR 3.5 (95% C.I. 1.8 – 7.3) for age <65 years; OR 5.1 (95% C.I. 2.7 – 10.2) for age ≥ 65 years. Overall, 70.4% (N=133) of cases and only 37.2% of controls (N=68) were found to have mutations. Mutations were found in 55.6% of cases and 26.0% of controls < 65 years old, and in 81.5% of cases and 45.3% of controls age ≥ 65 years.

**Table 1.**
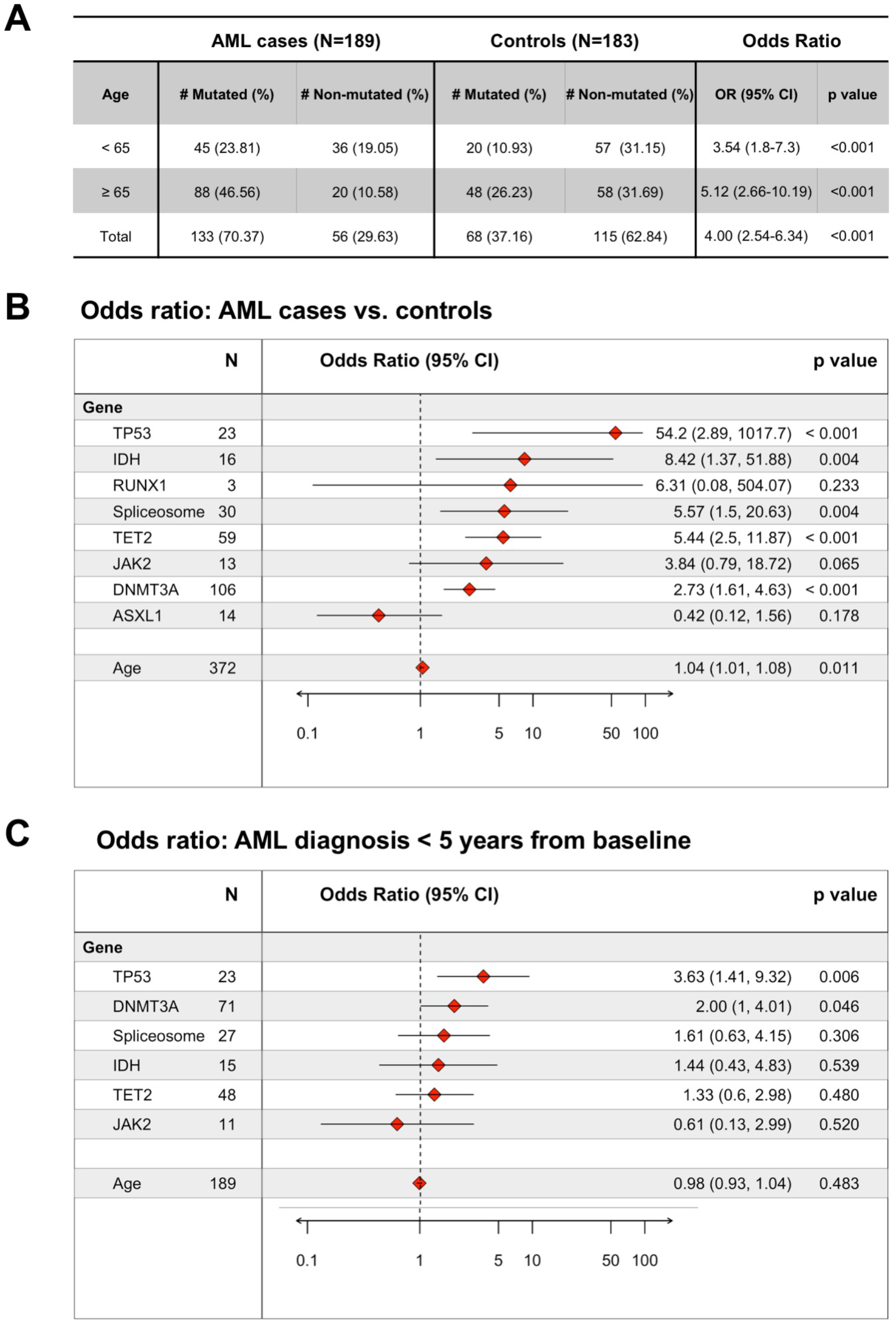
Mutation frequencies and odds ratios of AML. (A) Number and frequency of mutations in AML cases vs. controls overall and for participants younger than 65 years vs. ≥ 65 years. (B) Forest plot indicating odds ratio of mutations in each gene occurring in the AML cases vs. controls. Genes or gene categories significantly associated with AML cases include TP53, IDH, spliceosome, TET2, and DNMT3A. (C) Forest plot indicating odds ratios of AML diagnosis occurring sooner than 5 years. Mutations in TP53 and DNMT3A are significantly associated with rapid development of AML. IDH category includes IDH1 and IDH2. The spliceosome category includes SRSF2, SF3B1, and U2AF1. Abbreviations: CI, confidence interval; N, number affected. P-values are shown for penalized likelihood multivariable logistic regression.

Among the recurrently mutated genes, some mutations demonstrated increased specificity and penetrance for the development of AML. All participants with a TP53 (N=23/23) or RUNX1 (N=3/3) mutation developed AML. Also, all participants with an IDH1 or IDH2 mutation eventually developed AML except one control, which was lost to follow-up at the end of the study. Multivariable analysis was performed to evaluate potential associations between individual mutated genes and the development of AML, adjusting for confounders including co-mutated genes and age (Table 1B). This analysis found that TP53 (OR 54.2, 95% C.I. 2.9-1017.7), IDH (including IDH1 and IDH2) (OR 8.4, 95% C.I. 1.4-51.9), spliceosome genes (including SF3B1, SRSF2 and U2AF1) (OR 5.6, 95% C.I. 1.5-20.6), TET2 (OR 5.4, 95% C.I. 2.5 – 11.9) and DNMT3A (OR 2.7, 95% C.I. 1.6 – 4.6) were associated with significantly increased odds of developing AML relative to controls. The odds ratio was elevated for JAK2 (OR 3.8, 95% C.I. 0.8 – 18.7), but did not reach statistical significance. However, the specific JAK2 mutation JAK2 p.Val617Phe was significantly associated with AML development (OR 6.1, 95% C.I. 1.2 – 61.1, P = 0.03). Mutations in IDH1 and IDH2 were exclusively found in the known recurring hotspots in arginine-132 or arginine-140 (2, 13). Similarly, SRSF2 mutations were confined to the well-known proline-95 hotspot (14, 15). TP53 mutations were primarily in the DNA binding, transactivation, and oligomerization domains (16, 17). The distribution of mutations within functional protein domains, as well as the type of mutations found at each location, are shown in the Supplemental Data (Figures S3 – S6, S10 – S12).

### Mutations are associated with an accelerated time to AML presentation

AML cases with mutations at baseline experienced significantly shorter latency of disease than cases without baseline mutations. The presence of any mutation shifted the median time to AML diagnosis from 11.9 years (no mutations) to a median of 8 years after baseline assessment (P < 0.001, log rank test; Fig. 2A). Univariable analysis of mutated genes demonstrated that median time to AML varied according to mutated gene (Fig. 2B): DNMT3A (7.1 vs. 10.6 years), TP53 (4.9 vs. 10.2 years), spliceosome genes (7.4 vs. 10.0 years) and RUNX1 (1.5 vs. 9.6 years). The presence of RUNX1 mutations seemed to be associated with especially rapid development of AML within 2 years, but the number of participants in this subgroup was small (N=3). Clonal complexity also affected the latency of disease as patients with 1 mutation developed AML in 9.1 years, but those with 2 or more mutations developed AML in only 6.9 years (Fig 2C).

**Figure 2.**
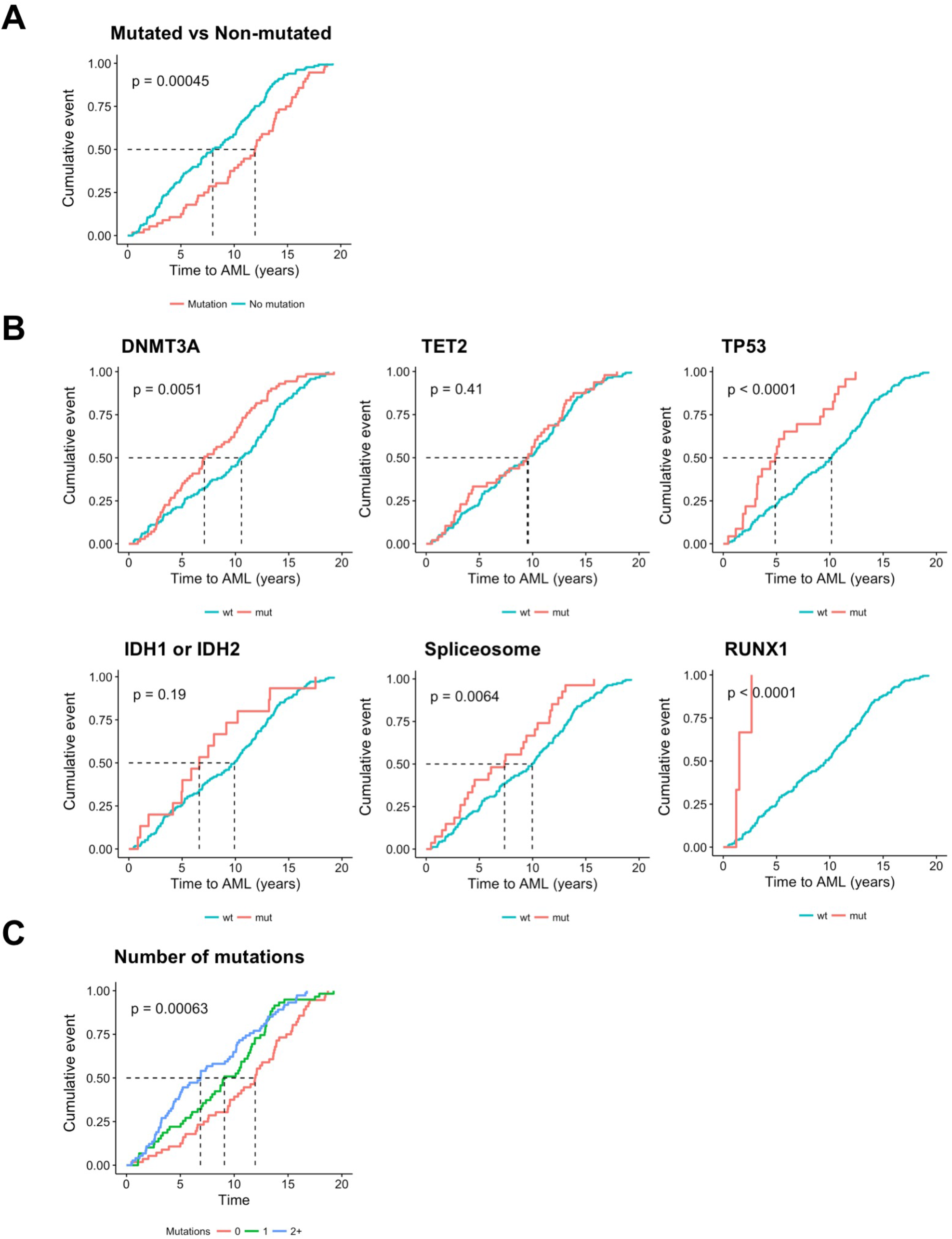
Time to AML diagnosis is influenced by mutation status. Cumulative incidence of AML diagnoses (cumulative event; vertical axis) as a function of time (years to AML diagnosis) is shown. Participants include AML cases only at baseline (N=189) with: (A) any mutated gene (Figure S4); (B) mutations in genes enriched in the AML cases and RUNX1; or (C) zero mutations, 1 mutations, or 2 or more (2+) mutations. P-values are shown following the log rank test.

Multivariable analyses produced similar findings (Table 1C). Mutations in TP53 were independently associated with AML occurring within 5 years of baseline (OR 3.6, 95% C.I. 1.4-9.1, P = 0.006) as well as an elevated annual risk of AML (HR 2.8, 95% C.I. 1.7-4.7, P < 0.001) when adjusted for confounding co-mutations and age. Similarly, DNMT3A mutations presented a smaller, but still significant risk of developing AML within 5 years (OR 2.9, 95% C.I. 1.0-4.1, P = 0.040), as well as an elevated annual risk (HR 1.6, 95% C.I. 1.2-2.2, P = 0.002). For TP53 and DNMT3A mutations, there was no appreciable difference in time to development of AML by mutation subtype, e.g. arginine-882 in DNMT3A. Finally, IDH mutations had an elevated annual risk of similar magnitude as DNMT3A (HR 1.6, 95% C.I. 0.9-2.8, P = 0.09), but this did not achieve statistical significance. However, while 15/16 participants with IDH mutations were eventually diagnosed with AML, there was no elevated risk of AML earlier than 5 years after baseline assessment, suggesting the possibility that further downstream mutational events are required for AML development.

### Mutations occurring at any variant allele fraction (VAF) in high-risk genes are associated with increased odds of AML

Of participants harboring any mutation evaluated at VAF > 10% at baseline, 83% eventually developed AML (OR 6.5, 95% C.I. 3.4 – 13.0, P < 0.001). When we considered only mutations in genes associated with development of AML shown in Fig. 3A, mutations present at VAF > 10% at baseline led to AML in 90% of cases (OR 11.6, 95% C.I. 5.1 – 31.1, P < 0.001). As demonstrated in the histograms, mutations in DNMT3A and TET2 at lower VAFs were notably more distributed among cases and controls. In contrast, mutations present at higher VAFs in DNMT3A and TET2 were almost exclusively seen in AML cases, suggesting that mutations in DNMT3A and TET2 are less specific for AML cases at lower VAFs. Thus, when excluding DNMT3A and TET2, 92% of patients harboring mutations at any VAF above the 1% cutoff (range 1.0%-54.9% VAF) in the remaining genes eventually developed AML (OR 16.5, 95% C.I. 6.4 – 54.0, P < 0.001). Next, we further examined the specificity mutations in genes significantly associated with AML cases (Table 1b) by determining their frequency in the controls. The true positive and false positive rate of mutations at varying VAF cutoffs was visualized using receiver operating characteristic (ROC) analysis (Fig. 3B). At a VAF cutoff of 1.3%, the presence of a mutation in any gene (excluding DNMT3A) resulted in a 4.9% rate of false positive detection (controls misclassified as cases). Individually, mutations in TP53, SRSF2, U2AF1, SF3B1, and IDH2 produced a ~1% rate of false positive detection at VAF cutoffs ranging from 1-2%. Tabulated results are available in Figure S7.

**Figure 3.**
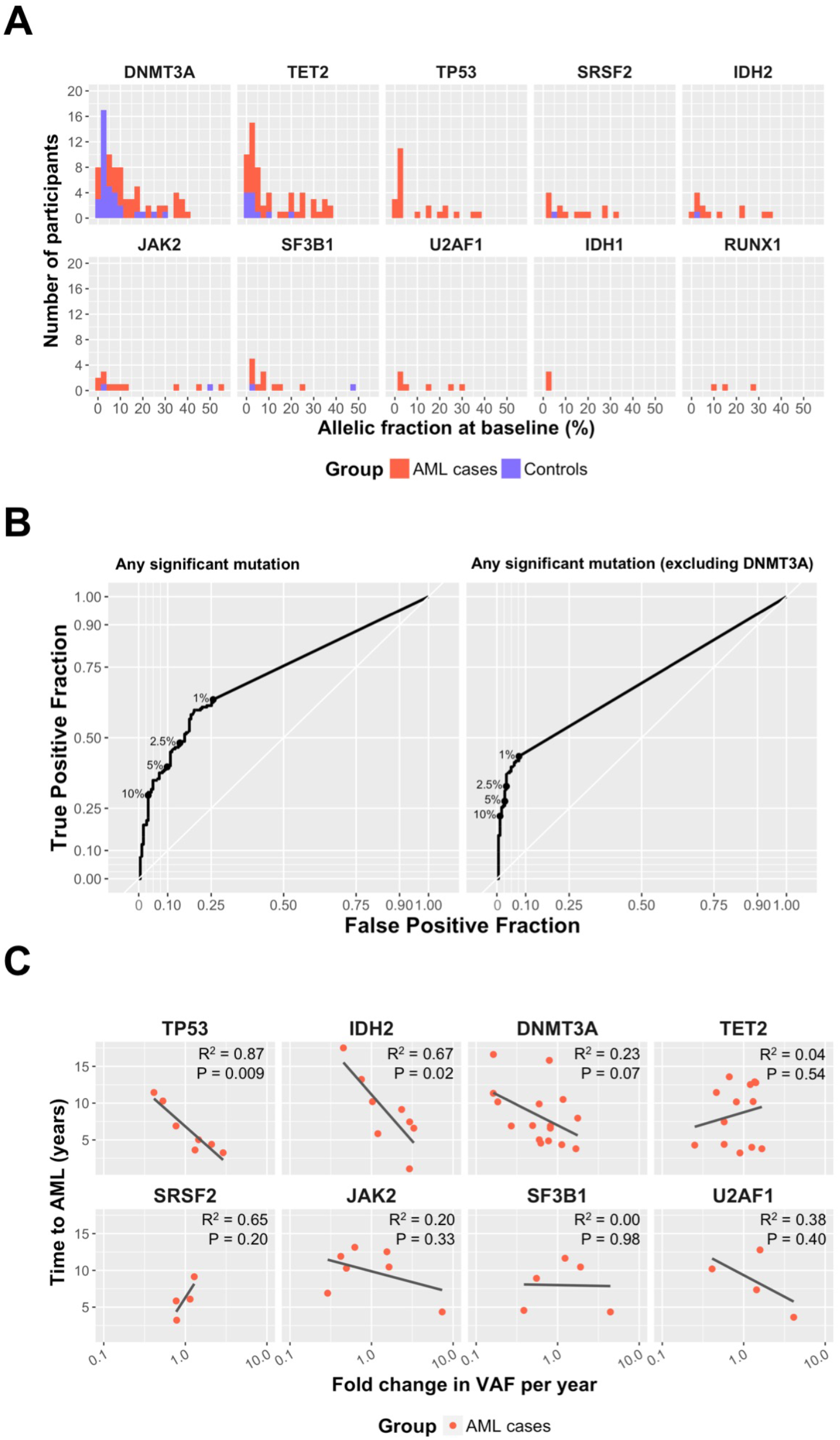
Mutations pose AML risk irrespective of the variant allele fraction. (A) Histogram indicating the maximum allelic fraction for mutations in each gene shown per participant at baseline. The proportion of AML cases (red) and controls (blue) is shown for each bin in the histogram. (B) Receiver operating characteristic (ROC) curves indicating the % true positive rate (vertical axis) vs. the % false positive rate (horizontal axis) of detecting AML cases. The curves indicate performance at each given allelic fraction (%) with 1%, 2.5%, 5%, and 10% indicated specifically (filled black circles). Performance is shown for mutations in any gene significantly associated with the AML case group (left plot; DNMT3A, TET2, IDH1, IDH2, SRSF2, SF3B1, U2AF1, TP53) or the same set of genes excluding DNMT3A (right plot). (C) Fold change in variant allele fraction (VAF) per year influences kinetics of AML diagnosis for TP53 and IDH2. Time to AML (years; vertical axis) is plotted against fold change in VAF as determined by comparing the VAF at baseline vs. the VAF at year 1 or year 3. Regression line is shown for each mutation in each gene. P-values shown reflect linear regression. Data on RUNX1 are provided since all participants with a RUNX1 mutation rapidly developed AML (< 2 years) although significance was not achieved due to the few participants mutated in RUNX1 within the cohort.

For the subset of participants for whom serial samples were available, changes in VAF were assessed from baseline to year 1 or year 3. We evaluated whether the time to AML development was influenced by the fold increase in VAF in participants who had demonstrated a statistically significant rise in VAF upon serial testing (Fig. 3C). The rate of increase in VAF for IDH2 mutations (R^2^ = 0.87, slope -3.3, 95% C.I. – 21.5 - -3.7, P = 0.02) and TP53 mutations (R^2^ = 0.67, slope -3.0, 95% C.I. -14.0 - -5.5, P = 0.009) was significantly associated with a shorter time to AML in a linear regression analysis. While this relationship was similar for mutations in DNMT3A, it did not achieve statistical significance. There were fewer changes from baseline in VAF across all genes in year 1, compared to year 3 (Figure S8). Finally, for participants with serial samples available, 89% of mutations detected in the study persisted from baseline to years 1 or 3, irrespective of whether the VAF significantly changed (Figure S9).

### Association between non-AML cancer history and mutations

We examined the mutational patterns of the 22 cases and 21 controls that had a history of malignancy at baseline, including breast, lung, bladder, endometrial, ovarian, colon cancer and lymphoma. Forty-five percent of AML cases (N=10/22) and 33% of controls (N=7/21) with a prior history of cancer at baseline were found to have mutations (not significant). While the absolute number of cases and controls with prior cancer history was few, we did note that 3/10 cases with prior history of cancer harbored IDH2 mutations at baseline evaluation and that all 3 of these cases had prior breast cancer. None of the cases or controls with a prior history of cancer had TP53 gene mutations.

### Progression to AML from baseline mutations

Despite having selected for participants who eventually developed AML, we noted the absence of NPM1 and FLT3 mutations, which are among the two most frequently recurring driver mutations in AML (1, 18). This finding is consistent with other reports of their absence in clonal hematopoiesis, and suggests that they may play a cooperative role in AML pathogenesis (4, 5). We identified a single case where follow up at year 1 preceded AML diagnosis by < 30 days (Fig. 4, Case A) and compared mutations at baseline and year 1 (< 30 days prior to AML diagnosis). The depth of sequencing coverage at NPM1 insertion sites was similar at both time points (~450x). The participant had an IDH2 mutation (8% VAF) at baseline. DNA from the sample acquired 1 year after baseline showed a new NPM1 type A mutation (14% VAF), along with the IDH2 mutation with increased VAF of 13%. AML was diagnosed less than 30 days later. The rapid development of AML after the acquisition of an NPM1 mutation suggests cooperation with the pre-existing IDH2 clone.

**Figure 4.**
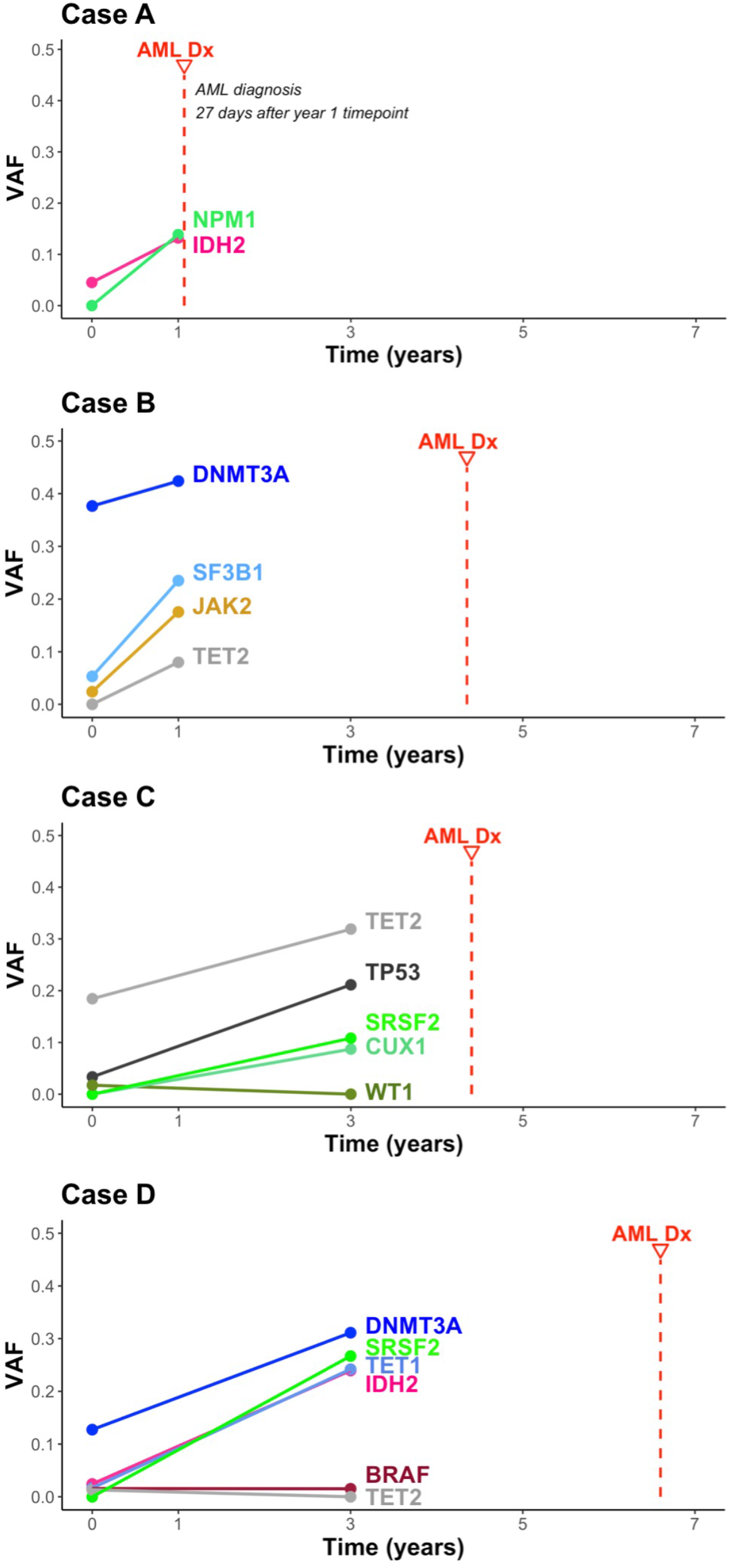
Clonal evolution towards AML in selected patients. Clonal composition and evolution are shown for four selected examples of participants who were evaluable serially (cases A, B, C, and D). Peripheral blood was sampled at baseline and years 1 or 2. The horizontal axis indicates time (years). The vertical axis indicates the VAF where the maximum possible VAF is 1 (100%). Mutated genes are shown at each time point as indicated on the line chart. Time of AML diagnosis (AML Dx) relative to baseline is indicated by the red vertical dotted line. **Case A**: An IDH2 mutation (8% VAF) is present at baseline at lower VAF and persists at year 1 at 13% VAF with an acquired NPM1 type A mutation at 14% VAF. AML diagnosis occurs < 30 days after the year 1 sample. **Case B:** DNMT3A mutation remained stable from baseline to year 1 follow up. VAFs of JAK2 and SF3B1 increased from 5% to 24% before AML diagnosis at 4.3 years from baseline. **Case C:** Clonal expansion of TP53 from 3% to 21% with acquired SRSF2 and CUX1 mutations between baseline and year 3 follow up. TET2 remains stable. AML diagnosed at 4.4 years from baseline. **Case D:** Low VAF mutations in IDH2 and TET1 expand by year 3 along with acquisition of SRSF2 in the presence of a relatively stable DNMT3A mutation. AML diagnosis occurs 6.6 years from baseline.

Other patterns of clonal evolution and expansion are demonstrated in representative cases B, C, and D. Mutations in genes typically associated with clonal hematopoiesis, such as DNMT3A and TET2, were shown to generally have stable or minimally increased VAFs in follow-up. Progression to AML was often preceded by the acquisition of new mutations, or by the expansion of mutations in other genes, such as RUNX1 or TP53 (Fig. 4).

## DISCUSSION

Peripheral blood samples collected as part of the WHI afforded a unique and unprecedented opportunity to investigate the acquisition of somatic mutations in AML patients prior to the development of overt disease. We found that study participants who eventually developed AML were significantly more likely to have mutations at WHI baseline evaluation and had higher mutational complexity than controls. The presence of a mutation at baseline was associated with increased odds of developing AML. Individually, the most significant mutations associated with increased odds of AML included those in TP53, IDH1/2, spliceosome (SRSF2, SF3B1, U2AF1), TET2, and DNMT3A. The median time to development of AML was shorter in subjects with mutations present at baseline WHI evaluation and those participants with baseline TP53 or DNMT3A mutations were more likely to develop AML within 5 years. Participants with mutations in the RUNX1 gene all developed AML within 2 years from baseline, but the numbers were too few to make statistical inferences. The time-to-AML was inversely correlated with increasing VAF in TP53, IDH2, and possibly DNMT3A mutations. While having a mutation at baseline increased the risk of developing AML, we identified a set of high-risk genes with especially high penetrance, including TP53, IDH2, SRSF2, SF3B1, and U2AF1. Any detectable VAF in the high-risk genes at a sensitivity cutoff of 1% was associated with increased risk of AML. When serial samples were available, we found that a rise in the allelic fraction of IDH2 or TP53 mutations was significantly associated with shorter time to AML in a manner directly proportional to the rate at which the allelic fraction increased. As expected, serial samples also revealed the stepwise acquisition of mutations leading to AML. Mutations in genes commonly associated with clonal hematopoiesis such as DNMT3A and TET2 were maintained over time, while new dominant sub-clones arose in genes such as NPM1, TP53 and SRSF2 preceded the development of AML.

Population-based studies reporting an association between CHIP mutations and increased risk of hematological malignancies have been described previously (4, 5). However, these populations included only a few cases of AML. In our large study, which included 189 AML cases, we found a significantly higher mutation frequency in both cases and controls (37% in cases vs. 10% in controls older than 65 years, and 23.4% in cases vs. 5.6 % in controls younger than 65 years). The higher mutation frequency in our study is most likely because we used a cutoff of 1% VAF with a high depth of coverage, while previous studies used a cutoff of 2% VAF or more. Also, our study population was somewhat older (median age 65 years) than in previous work and included only women. Older individuals are known to have more mutations and women in particular have been reported to have elevated incidence of subclonal DNMT3A mutations (19).

Age-related clonal hematopoiesis with TET2 mutations has been associated with slight neutropenia (20). In contrast, we did not find any statistically significant difference in baseline hematological parameters between cases and controls either overall, or by specific mutations. Our data do not suggest that routine CBC monitoring in a normal population will be a useful screening tool for AML, but it is possible that hematological parameters in our cohort changed during follow up from the baseline analysis as the mutant clones expanded. Thus, serial, prospective monitoring of standard hematological parameters in high-risk populations may be of value and should be investigated in conjunction with serial sequencing studies, as described below.

Two case-control studies of therapy-related AML and cancer controls have been published (21, 22). Gillis et al. analyzed 16 cases of therapy-related AML (t-AML) compared to 56 controls with chemotherapy exposure, but without AML (21). The authors concluded that t-AML cases were more likely to have a mutation compared to controls (OR 5.75, 95% CI 1.52-25.09, P = 0.01), but there was no difference in the prevalence of TP53 mutations among t-AML cases and non-AML cancer controls. In contrast, we found TP53 mutations exclusively in our AML cases. Also, none of our participants with a history of cancer prior to baseline assessment had TP53 mutations. One explanation for this observation could be that while the former study focused on t-AML cases only, our participants were from a normal population. It is possible that TP53 clones arising in response to chemotherapy might behave differently from clones arising de novo at VAFs exceeding 1% in healthy individuals. In a paper by Takahashi et al, 14 cases of therapy-related myeloid neoplasms and 54 cancer controls that did not develop myeloid neoplasms were analyzed for evidence of pre-existing clonal mutations (22). In this study, clonal hematopoiesis was detected in 71% of cases and 31% of cancer controls, which is close to the mutation frequency we detected in our cohort. Moreover, the study found TP53 mutations to be more frequent in therapy related myeloid malignancies, suggesting a slight clonal advantage driven by therapy-induced stress state.

A major strength of our study is having a cohort of normal controls that was reliably followed over 10 years for outcomes, including detailed baseline demographics, medical history, and laboratory data, thus providing a solid scientific base for a matched case-control analyses at the population level. A limitation of our study is that we could not perform sequencing studies of blood or bone marrow taken at the time of AML diagnosis. However, there were 4 cases in our study with serial sample collections drawn within a year from the diagnosis of AML and we did note persistence of previously detected mutations in the peripheral blood. Several commonly mutated genes in AML were absent in our study, notably FLT3 and NPM1, suggesting that the acquisition of these mutations may be later events in AML ontogeny. In our study, pre-existing TP53 and spliceosome gene mutations were strongly associated with the development of AML, but it is well-known that these mutations are not exclusive to AML and are also associated with other hematological malignancies, including myelodysplastic syndromes and chronic lymphocytic leukemia (23). Thus, further prospective monitoring studies will be required to confirm the specificity, or lack thereof, of these mutations in predicting AML versus other hematological disorders. In our study, we determined that for specific genes (i.e. TP53, IDH2, SF3B1, SRSF2, U2AF1), a VAF cutoff correlating with a false-positive fraction of less than 1% could be achieved. We propose the term “clonal hematopoiesis of AML potential (CHAP)” to be used for mutations associated with an increased risk of AML that can be detected at a VAF cutoff correlating with a false-positive fraction of less than 1% in controls.

In conclusion, our study demonstrates that detectable mutations likely arising from pre-leukemic clones are present in peripheral blood of individuals at a median of 9.8 years prior to the diagnosis of AML. The presence of mutations in TP53 or multiple mutations lead to especially rapid presentation of AML. This latency period suggests a possible window for therapeutic intervention prior to disease onset. To that end, we propose that patients who may be at increased risk for the development of AML should be followed longitudinally on long-term monitoring studies to further evaluate the prognostic and predictive importance of CHAP. Data from these studies will provide a robust rationale for clinical trials of preventative intervention strategies.

## ACKNOWELDGEMENTS

The authors would like to gratefully acknowledge all our donors to Leukemia Fighters, without whom this work would not have been possible. We would also like to acknowledge the women who generously participated in the WHI study. We are thankful to Dr. Jenny Z. Xiang and the Weill Cornell Genomics Core Facility as well as Jeffrey Catalano of the Englander Institute for Precision Medicine for assistance in sequencing. The WHI program is funded by the National Heart, Lung, and Blood Institute, National Institutes of Health, U.S. Department of Health and Human Services through contracts N01WH22110, 24152, 32100-2, 32105-6, 32108-9, 32111-13, 32115, 32118-32119, 32122, 42107-26, 42129-32, and 44221”, and the Cancer Center Support Grant NIH:NCI P30CA022453. We acknowledge the dedicated efforts of investigators and staff at the Women’s Health Initiative (WHI) clinical centers, the WHI Clinical Coordinating Center, and the National Heart, Lung and Blood program office (listing available at <u>http://www.whi.org</u>). We are additionally grateful for funding from the Sandra and Edward Meyer Cancer Center, which partially supported this study (D.C.H.).

